# Video-rate three-photon imaging in deep *Drosophila* brain based on a single Cr:forsterite oscillator

**DOI:** 10.1101/2023.03.23.533955

**Authors:** Lu-Ting Chou, Shao-Hsuan Wu, Hao-Hsuan Hung, Je-Chi Jang, Chung-Ming Chen, Ting-Chen Chang, Wei-Zhong Lin, Li-An Chu, Chi-Kuang Sun, Franz X. Kärtner, Anatoly A. Ivanov, Shi-Wei Chu, Shih-Hsuan Chia

## Abstract

We have demonstrated 30-Hz three-photon imaging using a single 24-MHz mode-locked Cr:forsterite oscillator with a center wavelength at 1260 nm. By managing the dispersion distribution in the resonator using double-chirped mirrors, we have produced 32-fs pulses with 22-nJ pulse energy. Using the oscillator as a driving source, we have realized multi-color three-photon images using a GFP-labeled *Drosophila* brain and an AF647-labeled mouse brain. To demonstrate the capability of deep-tissue imaging, we have obtained a 10-times higher SBR from the three-photon images than the two-photon results at different depths in a GFP-labeled *Drosophila* brain dissection. Furthermore, we have shown the impact of excitation pulse width on three-photon deep-tissue imaging. Our results indicate the superiority of using shorter pulses for deeper-tissue imaging, especially in the *Drosophila* brain. In addition, we have recorded the three-photon calcium imaging *in vivo* from the *Drosophila* mushroom body in response to external electric shocks. We believe our demonstration provides a robust approach for high-speed three-photon microscopy applications, especially for intravital investigations in the *Drosophila* brain.

## Introduction

The ability to visualize cellular structures and functions deep into biological tissues keeps improving with the advancement of optical technology.^1-3^ With the development of femtosecond lasers, multiphoton fluorescence microscopy has become a more critical tool for realizing deep-tissue structural and functional imaging at a sub-cellular resolution.^4-7^ Especially for brain studies, three-photon microscopy (3PM) has allowed imaging in deep brain regions that are typically inaccessible using single/two-photon microscopy, such as reaching the hippocampus of the intact mice brain8^1-3,8-10^ and realizing whole-brain observation using *Drosophila*.^11,12^ The reasons for the imaging depth extension using 3PM are two folds: Using femtosecond lasers in the transparent windows (*i*.*e*., around 1300 nm and 1700 nm) enables better tissue penetration of the excitation light with less scattering. (2) Employing higher-order nonlinear excitation improves the imaging contrast at depth by suppressing the out-of-focus background.

Despite the advantages of 3PM, the use of higher-order nonlinear excitation also implies a higher demand on the excitation source, and it is thus of great importance to discuss the limitation related to the source development. Compared with the 1700-nm transparency window, the 1300-nm excitation band has attracted more attention due to its compatibility with multi-color fluorescent molecules,^8,13^ including genetically encoded calcium indicators (*e*.*g*., GCaMPs). For generating energetic pulses centered around 1300 nm, the current state of the art relies on using optical parametric amplifiers (OPAs) pumped by bulky µJ-level solid-state sources at a repetition rate lower than 2 MHz.^2,3,9,10,14-16^ The lower repetition frequency limits the sampling rate using point scanning microscopes. For example, video-rate imaging is beneficial for *in vivo* observation, potentially allowing the investigation of fast bio-activities and preventing the imaging blur from unwanted movement;^10,14^ however, the 30-Hz frame rate with a typical size of 512×512 pixels requires the sampling rate at least 8 MHz. Although some works have demonstrated video-rate 3PM, such as using an adaptive excitation source,^3^ either the regions of interest are limited,^3^ or the pixel number over the whole field of view is compromised.^12,14^ As a result, the low pulse repetition rate limits the capability of video-rate imaging.

Besides the limitation on the repetition rate, the complexity and high cost of the OPA-based driving source hamper its general applicability to biomedical studies. Without additional amplification and nonlinear conversion, a long-cavity Cr:forsterite (Cr:F) oscillator may be a promising candidate to generate energetic femtosecond pulses around 1300 nm for driving high-speed 3PM. A typical Cr:F oscillator delivers 1260-nm pulses with an output power of hundreds of milliwatts and a repetition rate of around 80 MHz. The driving wavelength potentially allows simultaneous 3PM imaging of multi-color fluorophores;^13^ Using an extended cavity with a repetition rate around 15-30 MHz leads to the boost of the pulse energy up to tens of nJ^17,18^ while preserving the feasibility for high-speed imaging.

Although previous works have demonstrated 3PM based on a Cr:F oscillator,^6,19-21^ further investigations are required to prove its potential for deep-tissue imaging. These works relied on the use of bright biomarkers, such as SypHer3s,^6,19-21^ eosin,^22^ and Hoechst,^23^ with a long image acquisition time (*i*.*e*., 0.7 seconds to a few seconds). The compatibility with commonly used indicators, such as green fluorescent protein (GFP) and GCaMPs, and the possibilities of high-speed imaging have not been addressed. Furthermore, the imaging contrast was not strong enough to realize deeper-tissue imaging beyond the two-photon limit.

We start with the cavity engineering of a long-cavity Cr:forsterite oscillator to optimize the output energy and pulse width. Shorter pulse generation supported by a broader spectrum leads to higher peak power, potentially resulting in stronger multiphoton excitation.^24,25^ We employ a 24-MHz oscillator with a high pumping power, and we use double-chirped mirrors (DCMs)^26^ to provide broadband dispersion control. Moreover, we can easily manage the dispersion distribution inside the resonator in a compact fashion based on DCMs. The optimized oscillator delivers 32-fs pulses with a 70-nm bandwidth and 22-nJ pulse energy.

Although we cannot obtain the same energy level as commercial OPA systems, we have demonstrated our capability of 3PE excitations using commonly used indicators in brain imaging without noticeable photodamage. We have shown the improvement of the signal-to-background ratio (SBR) of 3PM GFP imaging using our source compared with the two-photon imaging using 920-nm light.^8,11,12^ In addition, we have measured the SBR at different imaging depths in the *Drosophila* and mouse brains under different excitation pulse widths while maintaining the peak power at the sample surface. A shorter pulse down to 35 fs leads to a more efficient three-photon GFP excitation at different depths, especially for *Drosophila* brain imaging. Furthermore, we also record the three-photon excited GCaMP7f signal from live *Drosophila* under a 30-Hz imaging rate with 512×512 pixels. These results indicated that our 24-MHz Cr:F oscillator can be a promising tool for high-speed 3PM brain imaging, especially using *Drosophila*.

## RESULTS AND DISCUSSION

### Optimization of Cr:F laser For Efficient Three-Photon Excitation

Proper Cr:F resonator design leads to the generation of energetic femtosecond pulses around 1260 nm. A schematic of the oscillator is shown in Fig. 1, and we employ a standard z-fold cavity with 150-mm radius-of-curvature focusing mirrors. We use a Yb:fiber laser (YLM-20-LP, IPG) as the pumping source. A lens focuses the pump light with a focal length of 100 mm into a Cr:F crystal, and the diameter of the focused spot in the crystal is around 67 µm. We stabilize the temperature of the laser crystal at 278K with a liquid chiller and a thermoelectric (TE) cooler. The crystal is 13-mm long with an absorption constant of 0.5 cm-1. We use a low-absorption Cr:F crystal to relieve the thermal loading of the laser crystal, and thus we may obtain a high power output by using a high pump power of 14 W.^27,28^ Furthermore, we try to increase the pulse energy by expanding the cavity length with a lower repetition rate: We implement a concave mirror with a 2000-mm radius of curvature to relay the beam propagation in the cavity arm with a 10% output coupler, and the total cavity length is 6.22 meters.

**Figure 1.**
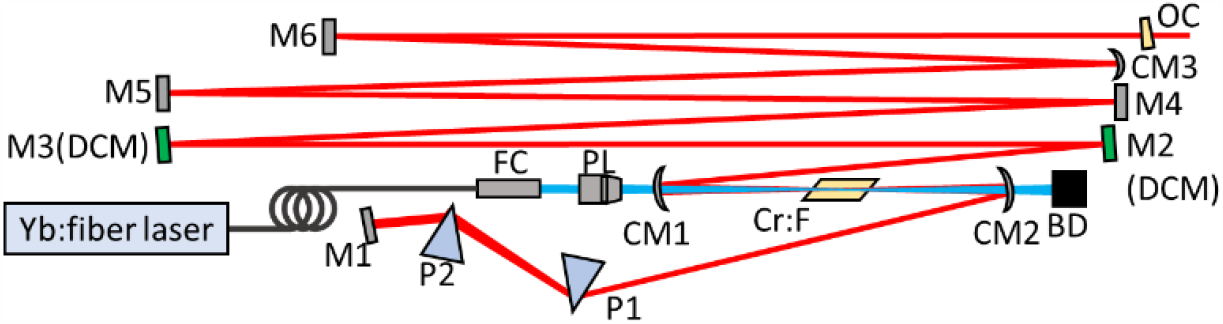
Schematic of the home-built mode-locked 24MHz Cr:F laser. FC: Fiber collimator; PL: Pump lens; CM: Concave mirror; DCM: Double-chirped mirror; M: Mirror; PL: Pump lens; P1 and P2: Prism 1 and 2; OC: Output coupler; BD: Beam dump.

Besides the pulse energy, we aim to generate shorter pulses with a broader spectrum from a Cr:F oscillator. Since the operating wavelength is close to the zero-dispersion wavelength of the crystal, higher-order dispersion becomes the main factor limiting pulse widths. A typical femtosecond Cr:F oscillator delivers sub-100-fs pulses, and the intracavity pulse shaping relies on the balance between self-phase modulation and negative dispersion in separate cavity elements over a resonator round trip. The mode-locking mechanism is similar to soliton formation, and it can be stable enough if the experienced phase shift per round trip is not too strong. Here, we manage the dispersion distribution in the resonator to optimize the output bandwidth and pulse width. We use a pair of SF14 prisms in the short arm of the linear laser cavity and optionally replace M2 and M3 with DCMs, as shown in Fig. 1. The DCM introduces -75-fs2 group delay dispersion per bounce around 1260 nm, and it offers broadband dispersion compensation with high reflectivity. These DCMs have been applied for the generation of 14-fs pulses with a low 80-mW average power.^26^ We experimentally compare the output performances under different dispersion balance schemes: (1) M2 and M3 in Fig. 1 are regular dielectric mirrors; (2) M2 is a DCM, and M3 is a regular dielectric mirror; (3) M2 and M3 are DCMs.

We tune the intracavity dispersion by changing the material insertion of the prism pair in each scheme. In Fig. 2(a), the transform-limited (TL) output pulse width varies in proportion to the net negative group delay dispersion. Introducing DCMs and realizing more balanced dispersion management can achieve stable mode-locking with less net intracavity dispersion with broader spectra and shorter output pulse widths. An output spectrum, which corresponds to the 30-fs TL width, is shown in Fig. 2(b) with a full width of half maximum (FWHM) bandwidth of 70-nm, and the average power is 530 mW, corresponding to a pulse energy of 22 nJ. We also characterize the pulse width using a home-built autocorrelator, and the direct output features around 32-fs pulse width under the assumption of a hyperbolic secant shape, as shown in Fig. 2(c). We have confirmed the operation of solitary mode-locking based on the linear tendency in Fig. 2(a) and almost-chirp-free performance at the direct output, as shown in Fig. 2(c). We want to note that the achievable peak power is up to 0.6 MW, which is twice the upper limit discussed in previous work.^18^

**Figure. 2.**
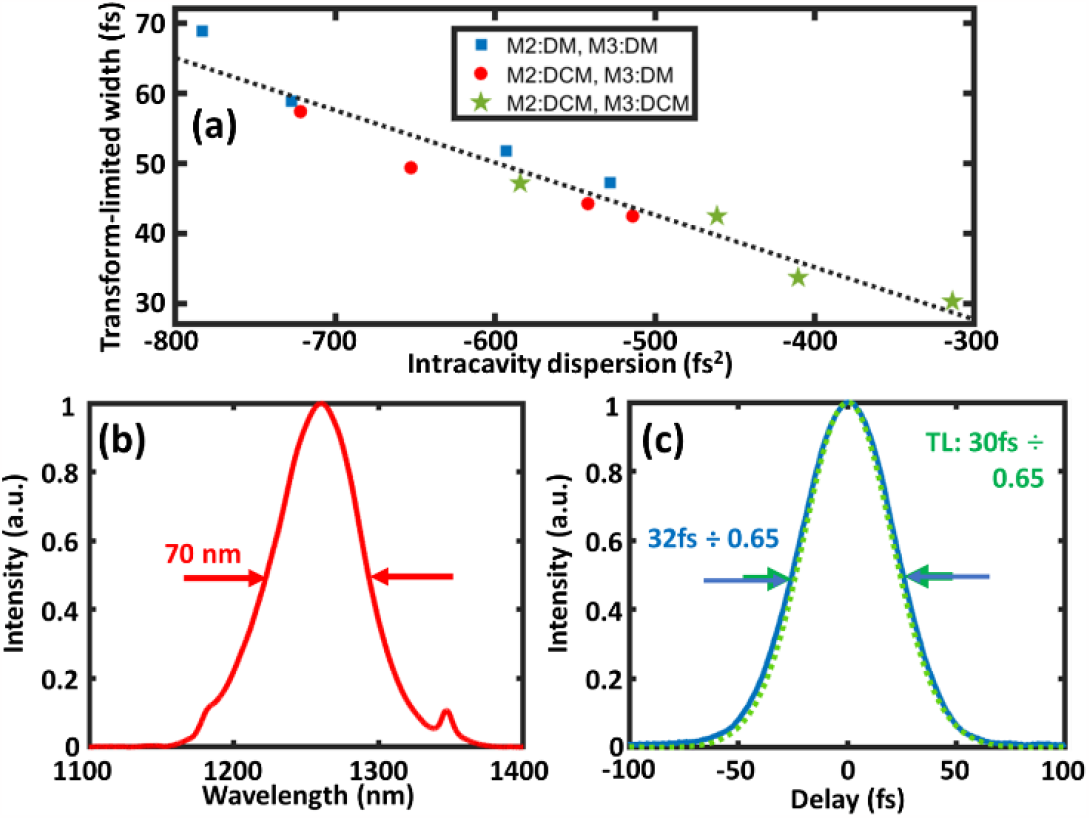
Output specification of the 24-MHz Cr:F laser. (a) TL pulse widths operated under different net intracavity GDD with the specified schemes. Blue square: M2 and M3 in Fig. 1 are regular DMs; red circle: M2 is a DCM, and M3 is a regular DM; green star: M2 and M3 are DCMs. All the data are close to a linear line (black dotted). (b) The optimized output spectrum. (c) The measured autocorrelation trace (blue solid) and the TL trace (green dotted). TL: Transform-limited; DM: Dielectric mirror; DCM: Double-chirped mirror; GDD: Group delay dispersion.

### Video-rate Green Fluorescent Three-photon Microscopy

To demonstrate the applicability of our 24-MHz sources for 3PM imaging, we use a home-built scanning microscope with a customized processing and acquisition unit (Jade Bio, SouthPort Co.) and a 40X 1.15NA objective lens (CFI Apo LWD Lambda S 40XC WI, Nikon). We separate the epi-detected fluorescence from the excitation by a longpass dichroic beam splitter (FF735-Di02, Semrock), and the signal is detected by a PMT (H7422, Hamamatsu) after the spectral isolation using an infrared blocker (700FESH, Thorlabs) and a bandpass filter (FF01-525/39 and FF01-660/25, Semrock, for the green and red fluorescence respectively). The output of the PMT is amplified by a transimpedance amplifier (TIA60, Thorlabs) and sent to the acquisition unit. To optimize the nonlinear excitation, we compress the illuminating pulses after the objective lens by a DCM pair before the microscopy system. We acquire all the images under a 250µm×250µm field of view (FOV) with 512×512 pixels and a 30-Hz frame rate. We want to note that no frame averaging is applied, and a higher imaging rate at 60 Hz is obtainable with 256×256 pixels.

First, the direct use of our Cr:F oscillator potentially allows the realization of multi-color 3PM.^13^ We employ two samples with distinct emission spectra, a GFP-expressed *Drosophila* brain dissection and a mouse brain sample slide with AF647-labeled neurons (FluoTissue mouse brain section, SunJin Lab). We have obtained clear GFP and AF647 images, as shown in Figs. 3(a) and (c), and we analyze the dependencies of the signal intensity on the illuminating power, as shown in Figs. 3(b) and (d), respectively. The signal strength in both cases is proportional to the cube of the excitation power. As a result, we have successfully demonstrated clear 3PM brain images using commonly used green and red indicators.

**Figure 3.**
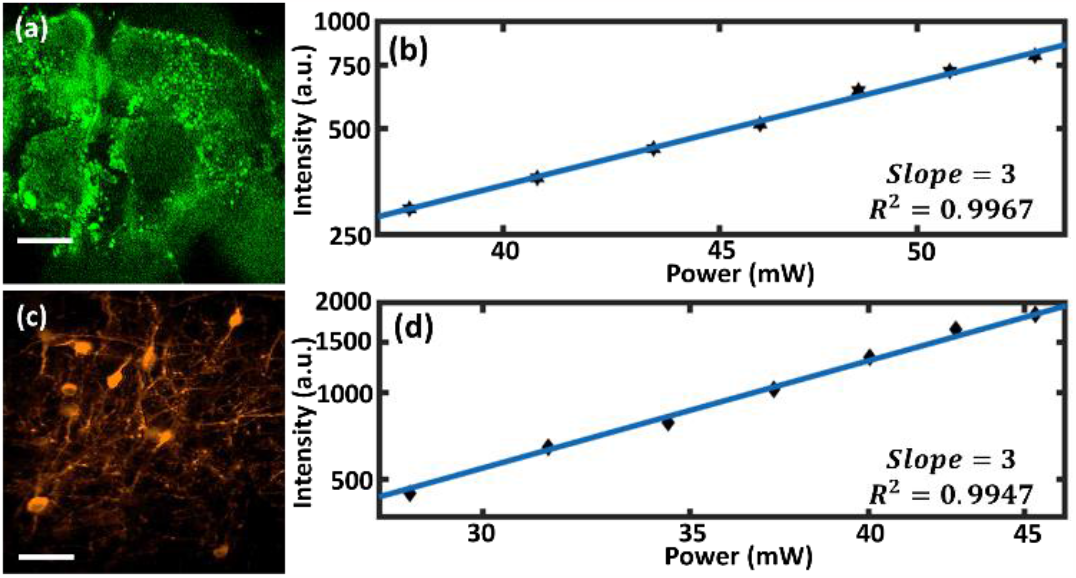
Demonstration of three-photon excitation using Cr:F laser. (a) Three-photon image of GFP-expressed *Drosophila* brain. (b) Cubic dependence of GFP signal in (a) on the excitation power. (c) Three-photon image of AF647-labeled neurons in a mouse brain slide. (d) Cubic dependence of AF647 signals in (c) on the excitation power. The power is measured after the objective lens. The scales of both axes in (b) and (d) are in the logarithmic scale. The scale bar is 50 μm.

It is interesting to see the cubic dependency of AF647 excitation. It is well known that AF647 exhibits a main linear absorption peak at 633 nm and a secondary one around 594 nm,^29^ but there is no report on the signal dependency of AF-647 using lasers around 1260 nm. Since the excitation wavelength of our laser is close to twice the wavelength of the one-photon absorption peak of AF647, the fluorescence signal can be excited by a mixture of two-photon and three-photon processes. Previous work has discussed the signal dependency on the pumping power at different wavelengths using Texas Red:^13^ When the two-photon and three-photon excitation cross sections are overlapped, using longer and shorter excitation wavelengths leads to more three-photon and two-photon transitions, respectively. The cubic power dependency in our measurement shows the dominance of the three-photon transition process, led by the excitation to the higher-energy bands. Moreover, we have used another 1150-nm laser^5^ to study the power dependency, and the result exhibits a slope of 2 in the logarithmic scale. The results indicate that the 594-nm peak is efficiently excited via two-photon absorption at 1150 nm, but the 633-nm peak is not by the 1260-nm excitation wavelength. The underlying photophysical mechanism and selection rule is interesting and requires further investigation.

We use the same GFP-labeled *Drosophila* brain dissection with a thickness of 150 µm to show our capability for deep-tissue imaging: We compare the 3PM images at different depths, as shown in Figs. 4(a)-(d), with the two-photon excitation ones, as shown in Figs. 4(e)-(h). We obtain the two-photon images using a 920-nm fiber source^5^ with a proper dispersion pre-compensation using a DCM pair (DCM7, Laser Quantum). To maximize the two-photon signal, we increase the illuminating power of the 920-nm light to 40 mW at the sample surface, and we may observe photodamage when applying a higher excitation power. It is worth noting that even though the brain thickness of *Drosophila* is only around 200-μm, common femtosecond sources for GFP two-photon excitation cannot penetrate, possibly due to light aberration/scattering of the trachea.^11,12^ On the other hand, using our Cr: F oscillator, we can illuminate a high average power of 100 mW on the sample surface without noticeable photodamage. We thus obtain clearer deep-tissue 3PM GFP images using our source, and the insufficient contrast from the two-photon imaging at depth would lead to difficulties in analyzing structural information.

**Figure 4.**
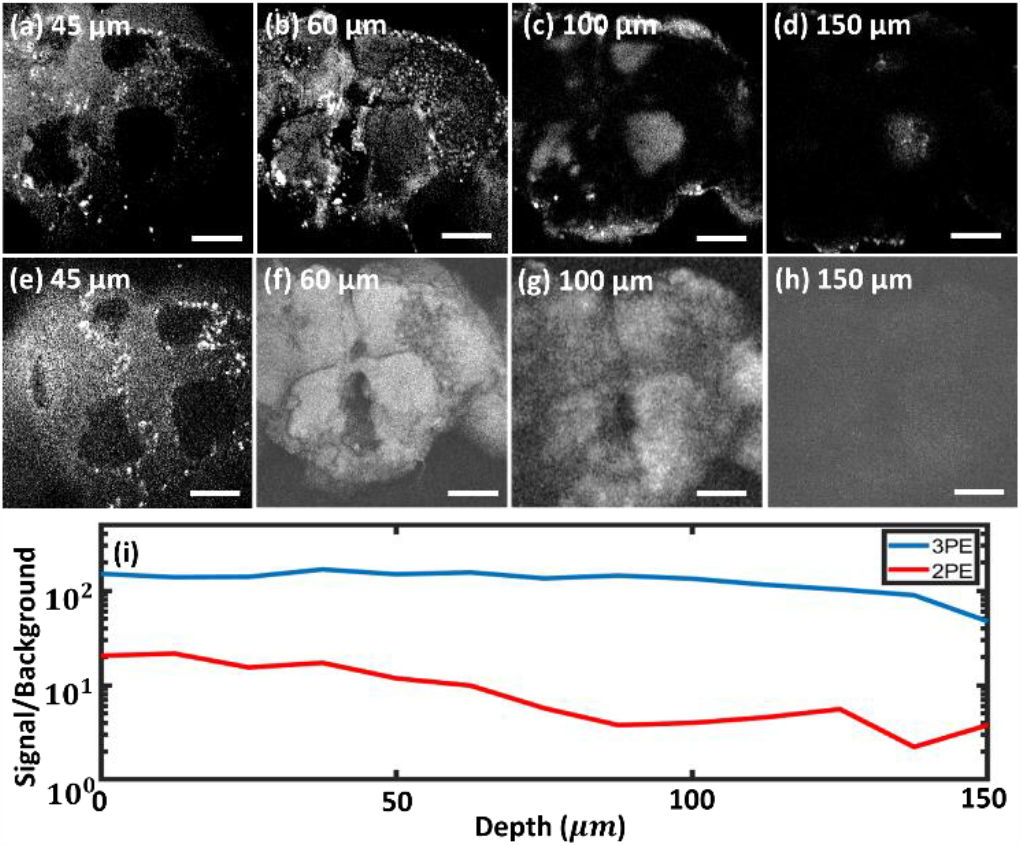
Comparison of SBR between 3PE and 2PE images. (a)-(d) 3PE images and (e)-(h) 2PE images in a GFP-labeled *Drosophila* brain at different imaging depths. (i) Comparison of SBR using 2PE (red) and 3PE (blue) at different imaging depths, and the vertical axis is on a logarithmic scale. The pixel size is 512×512, and the scale bar is 50 μm. 3PE: Three-photon excitation; 2PE: Two-photon excitation; SBR: Signal-to-background ratio.

We also calculate the SBR according to the previous work,^11^ as shown in Fig. 4(i). The SBR at different depths using the three-photon excitation is nearly ten times higher than the two-photon one. We have obtained a similar three-photon SBR as the previous work.^11^

We can easily tune the output pulse width and bandwidth of the Cr:F oscillator by changing the material insertion of the intracavity prism pair, and thus we may discuss the influence of the laser pulse width on the penetration depth and SBR of 3PM imaging. We have compared the SBR at different imaging depths using GFP-expressed *Drosophila* and mouse brain dissections, as shown in Figs. 5(a) and (b), respectively. We employ a DCM pair and a fused silica wedge pair to optimize the signal strength at the sample surface in each case, and we may obtain well-compressed pulses at the sample surface. We try to maintain the second harmonic generation signal from a thin BBO crystal after the objective lens by controlling the illuminating pulse energy, roughly proportional to the laser bandwidth and inversely proportional to the compressed pulse width. For example, we apply 2.7-nJ pulse energy when using 35-fs pulses with a spectral bandwidth of 60 nm, and 7.8-nJ pulse energy is used for 105-fs pulses with a spectral bandwidth of 20 nm. In Fig. 5(a), the shorter pulses lead to slightly lower, less than 10%, SBR near the *Drosophila* brain surface, possibly because of the more severe chromatic aberration of the scanning microscope system and the specimens with a broader excitation spectrum. However, the SBR led by shorter pulses becomes significantly higher in the deeper positions. At the imaging depth of 144 μm, the 35-fs pulses contribute to a three-times-higher SBR than using 105-fs pulses. We also apply the same protocol for the SBR comparison in a mouse brain sample labeled by GFP, as shown in Fig. 5(b). In the mouse brain, we can also observe a similar tendency that the SBR using shorter pulses is slightly lower near the surface and getting higher in deeper positions than using longer pulses. However, although we may obtain a deeper penetration depth of 600 μm in a mouse brain, the improvement of SBR at depth with shorter pulses is much less evident than that using the *Drosophila* brain dissection. We have thus shown that the shorter excitation pulses enable the higher SBR in the deeper layers,^30^ especially in the *Drosophila* brain with more severe light scattering.

**Figure 5.**
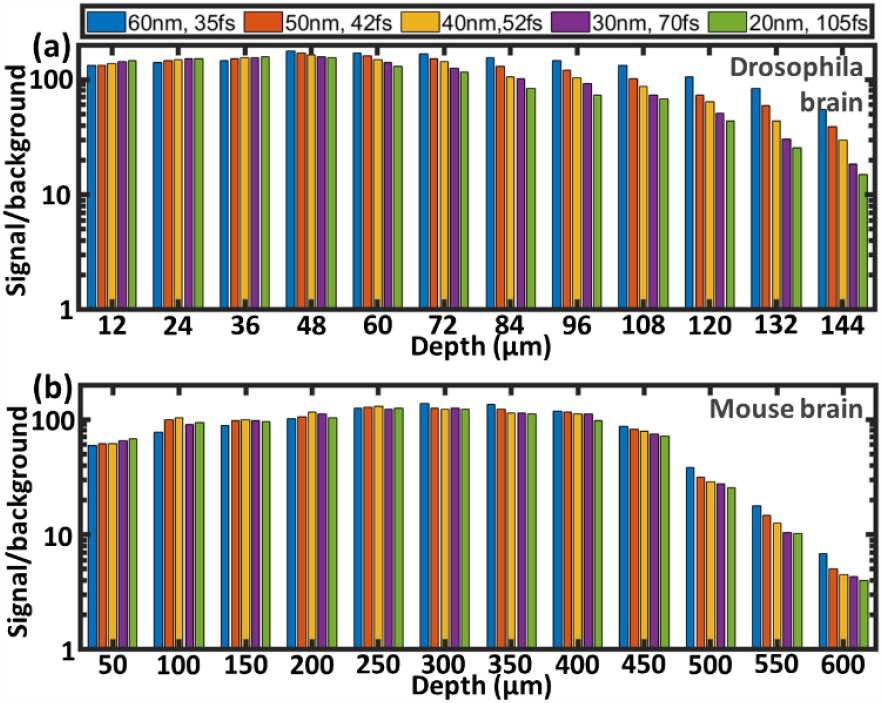
Signal-to-background ratio of 3PM imaging in different depths and tissues. SBR in (a) a GFP-labeled *Drosophila* brain and (b) a GFP-labeled mouse brain, under a fixed peak intensity. We use compressed pulses with specified bandwidths ranging from 60 nm to 20 nm, corresponding to the transform-limited pulse widths from 35 fs to 105 fs.

The demonstrated source aims for 30-Hz 3PM imaging for the structural and functional observation of physiological dynamics. Therefore, we test the ability of our laser for video-rate imaging by continuously scanning different depths in the *Drosophila* brain, and the screenshots have been shown in Figs. 4(a)-(d), where the video, labeled with the recorded depth and time, is available in Supplement 1 (https://reurl.cc/dejogk). We illuminate a high average power of 100 mW on the sample surface without noticeable photodamage, which is consistent with previous work.^31^ We also implement this source for *in vivo* video-rate monitoring of GCaMP7f signals expressed in the Kenyon cells of the mushroom body, which are important for revealing the learning and memory mechanisms,^32,33^ and we show the result in Fig. 6. GCaMP7f is commonly used to image the neuronal activity by tracking changes in calcium ions levels over precise timescales.^34^ We apply electric shocks to a live *Drosophila* while recording the images of a target area, as shown in the inset in Fig. 6. We slightly zoom in the target area to observe the details of the field of interest, and the field of view is 200μm×200μm with 512×512 pixels. We fix the live *Drosophila* on a metal plate with a hole, and we contact the belly of *Drosophila* with a wire covered by conductive gel so that the electric current can pass through the whole body of the *Drosophila*. We apply electric shock every 10 seconds using rectangular waves with a voltage of 15V via a function generator. The stimulation leads to the transfer of neural signals, which is associated with a change in concentration of calcium ions, and the released calcium ions combine with GCaMP7f for enhanced green fluorescence emission. We record and plot the GCaMP7f signal intensity along with time, and we label the stimulated periods as the red shallowed regions in Fig. 6. The fluorescence signal increases by around 15% after the stimulations, and we also plot the signal fluctuation when observing a rest *Drosophila* brain for reference. Moreover, we record the video-rate depth scanning in the GFP-labeled mouse brain, with 250μm×250μm field of view and 512×512 pixel number, which enables observing a large number of neurons with a high imaging speed, as shown in Supplement 2 (https://reurl.cc/zrjkLp). We also plot the intensity change at different positions during the depth scan, as shown in Fig. 7.

**Figure 6.**
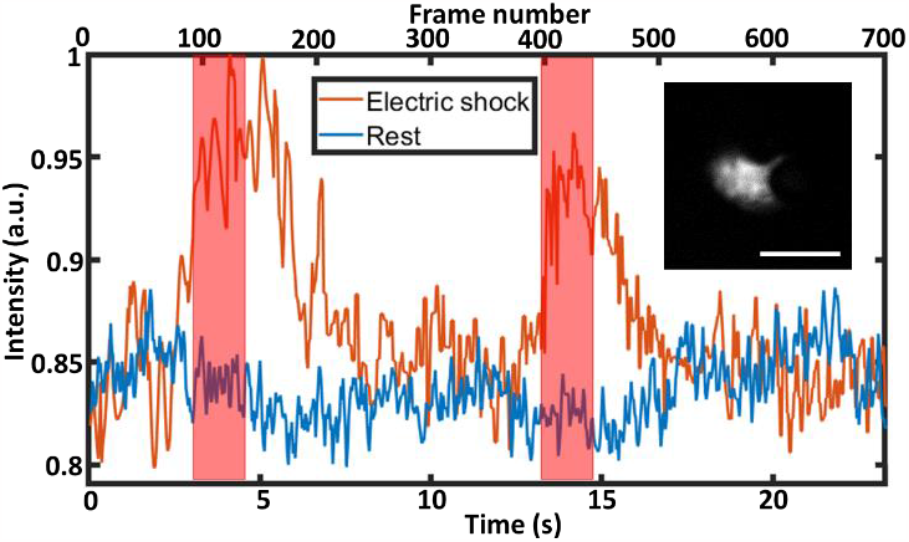
Video-rate monitoring of neuroactivity signal in a live *Drosophila* brain. Three-photon signal intensity of GCaMP7f from the mushroom body in a Drosophila brain in vivo under electric stimulation, and the signals are collected from the inset image. The red shallowed regions indicate the periods applying electric stimulation. The orange and blue lines indicate the GCaMP7f intensity with electric shocks and at rest, respectively, and the scale bar is 80 μm.

**Figure 7.**
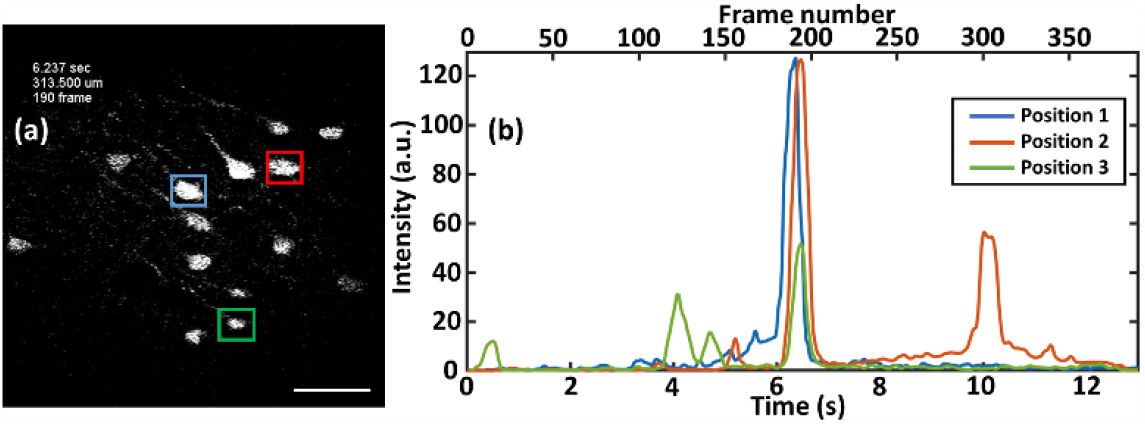
Video-rate recording of Drosophila brain at different depths. (a) A screenshot of video-rate depth scanning in the GFP-labeled mouse brain. The scale bar is 50 μm. (b) The fluorescent intensity at different positions (*i*.*e*., blue, red, and green frames as Position 1, 2, and 3 in (a)) and time. The scanning speed is 50 μm per second.

## CONCLUSION

We have demonstrated an optimized femtosecond source enabling high-speed 3PM, especially for deep *Drosophila* brain imaging. We build a Cr:F oscillator around 1260 nm for multi-color three-photon imaging, and we boost the output pulse energy by increasing the cavity length and relieving the thermal loading of the laser crystal under a high pumping power. Furthermore, we investigate the dispersion distribution in solitary mode-locking, and we have obtained shorter pulses with broader spectra by balancing the dispersion distribution inside the resonator. We have delivered 32-fs pulses with 22-nJ pulse energy from the direct oscillator output.

We also discuss the strengths in three-photon brain imaging using our Cr:forsterite oscillator. We have realized multi-color 3PM images using a GFP-labeled *Drosophila* brain and an AF647-labeled mouse brain. Moreover, we have obtained a 10-times higher SBR from 3PM images than the two-photon results at different depths in a GFP-labeled *Drosophila* brain dissection. Furthermore, we have investigated the impact of excitation pulse width on the SBR of 3PM at different imaging depths. We compare the SBR using *Drosophila* and mouse brains under a consistent peak power. The results indicate the superiority of using shorter pulses for deep-tissue imaging, especially in the *Drosophila* brain.

We have also demonstrated the 30-Hz imaging rate by continuously scanning through a 150-µm-thick *Drosophila* brain using 32-fs pulses. In addition, we record the neural signals *in vivo* from the *Drosophila* mushroom body in response to external electric shocks. Our source development potentially allows monitoring of rapid physiological dynamics in the deep tissue with a video rate. Although the output pulse energy is much smaller than commercial OPA systems, we believe our demonstration provides a robust approach for high-speed 3PM applications, especially for intravital investigations in the *Drosophila* brains.

## ACKNOWLEDGMENTS

This work was financially supported by the Young Scholar Fellowship Program from the National Science and Technology Council (NSTC) in Taiwan, under Grant NSTC 111-2636-E-010-002 (Shih-Hsuan Chia), and NSTC-111-2321-B-002-016 (Shi-Wei Chu).

## AUTHOR CONTRIBUTIONS

L.-T. C. and S. -H. C. conceived the study. L.-T. C., S. -H. W., and H. -H. H. built the Cr:F oscillator. L.-T. C. and W. -Z. L. Built the imaging system. L.-T. C., S. -H. W., and J. -C. J. performed imaging experiments. S. -H. W., C. -M. C., and T. -C. C. prepared the samples. L. -A. C. provided samples. C. -K. S provided supports and elements for Cr:F oscillator. F. X. K. provided supports and elements for pulse compression. L.-T. C., S. -H. W., and H. -H. H. processed and analyzed the data. A. A. I. provided supports for Cr:F oscillator construction and cavity design. S. -W. C. provided protocol and process for three-photon imaging of *Drosophila*. L.-T. C. and S. -H. C. wrote the article. S. -H. C. supervised the whole study.

## REFERENCES

1. Horton, N.G., Wang, K., Kobat, D., Clark, C.G., Wise, F.W., Schaffer, C.B., and Xu, C. (2013). In vivo three-photon microscopy of subcortical structures within an intact mouse brain. Nat. Photonics 7, 205–209.

2. Chow, D.M., Sinefeld, D., Kolkman, K.E., Ouzounov, D.G., Akbari, N., Tatarsky, R., Bass, A., Xu, C., and Fetcho, J.R. (2020). Deep three-photon imaging of the brain in intact adult zebrafish. Nat. Methods 17, 605–608.

3. Li, B., Wu, C., Wang, M., Charan, K., and Xu, C. (2020). An adaptive excitation source for high-speed multiphoton microscopy. Nat. Methods 17, 163–166.

4. Xu, C., and Wise, F. (2013). Recent advances in fibre lasers for nonlinear microscopy. Nat. Photonics 7, 875–882.

5. Chou, L.-T., Liu, Y.-C., Zhong, D.-L., Lin, W.-Z., Hung, H.-H., Chan, C.-J., Chen, Z.-P., and Chia, S.-H. (2021). Low noise, self-phase-modulation-enabled femtosecond fiber sources tunable in 740-1236 nm for wide two-photon fluorescence microscopy applications. Biomed. Opt. Express 12, 2888–2901.

6. Lanin, A.A., Chebotarev, A.S., Pochechuev, M.S., Kelmanson, I.V., Kotova, D.A., Bilan, D.S., Ermakova, Y.G., Fedotov, A.B., Ivanov, A.A., and Belousov, V.V. (2020). Two-and three-photon absorption cross-section characterization for high-brightness, cell-specific multiphoton fluorescence brain imaging. J. Biophotonics 13, e201900243.

7. Horton, N.G., Wang, K., Kobat, D., Clark, C.G., Wise, F.W., Schaffer, C.B., and Xu, C. (2013). In vivo three-photon microscopy of subcortical structures within an intact mouse brain. Nat. Photonics 7, 205.

8. Wang, T., and Xu, C. (2020). Three-photon neuronal imaging in deep mouse brain. Optica 7, 947–960.

9. Wang, T., Wu, C., Ouzounov, D.G., Gu, W., Xia, F., Kim, M., Yang, X., Warden, M.R., and Xu, C. (2020). Quantitative analysis of 1300-nm three-photon calcium imaging in the mouse brain. Elife 9, e53205.

10. Yildirim, M., Sugihara, H., So, P.T., and Sur, M. (2019). Functional imaging of visual cortical layers and subplate in awake mice with optimized three-photon microscopy. Nat. Commun. 10, 1–12.

11. Hsu, K.-J., Lin, Y.-Y., Chiang, A.-S., and Chu, S.-W. (2019). Optical properties of adult Drosophila brains in one-, two-, and three-photon microscopy. Biomed. Opt. Express 10, 1627–1637.

12. Aragon, M.J., Mok, A.T., Shea, J., Wang, M., Kim, H., Barkdull, N., Xu, C., and Yapici, N. (2022). Multiphoton imaging of neural structure and activity in Drosophila through the oikjuintact cuticle. Elife 11, e69094.

13. Hontani, Y., Xia, F., and Xu, C. (2021). Multicolor three-photon fluorescence imaging with single-wavelength excitation deep in mouse brain. Sci. Adv. 7, eabf3531.

14. Klioutchnikov, A., Wallace, D.J., Frosz, M.H., Zeltner, R., Sawinski, J., Pawlak, V., Voit, K.-M., Russell, P.S.J., and Kerr, J.N. (2020). Three-photon head-mounted microscope for imaging deep cortical layers in freely moving rats. Nat. Methods 17, 509–513.

15. Ouzounov, D.G., Wang, T., Wang, M., Feng, D.D., Horton, N.G., Cruz-Hernández, J.C., Cheng, Y.-T., Reimer, J., Tolias, A.S., and Nishimura, N. (2017). In vivo three-photon imaging of activity of GCaMP6-labeled neurons deep in intact mouse brain. Nat. Methods 14, 388–390.

16. Wang, T., Ouzounov, D.G., Wu, C., Horton, N.G., Zhang, B., Wu, C.-H., Zhang, Y., Schnitzer, M.J., and Xu, C. (2018). Three-photon imaging of mouse brain structure and function through the intact skull. Nat. Methods 15, 789–792.

17. Ivanov, A., Voronin, A., Lanin, A., Sidorov-Biryukov, D., Fedotov, A., and Zheltikov, A. (2014). Pulse-width-tunable 0.7 W mode-locked Cr: forsterite laser. Opt. Lett. 39, 205–208.

18. Ivanov, A.A., Martynov, G.N., Lanin, A.A., Fedotov, A.B., and Zheltikov, A.M. (2020). High-energy self-mode-locked Cr: forsterite laser near the soliton blowup threshold. Opt. Lett. 45, 1890–1893.

19. Pochechuev, M., Lanin, A., Kelmanson, I., Chebotarev, A., Fetisova, E., Bilan, D., Shevchenko, E., Ivanov, A., Fedotov, A., and Belousov, V. (2021). Multimodal nonlinear-optical imaging of nucleoli. Opt. Lett. 46, 3608–3611.

20. Lanin, A., Pochechuev, M., Chebotarev, A., Kelmanson, I., Bilan, D., Kotova, D., Tarabykin, V., Ivanov, A., Fedotov, A., and Belousov, V. (2020). Cell-specific three-photon-fluorescence brain imaging: neurons, astrocytes, and gliovascular interfaces. Opt. Lett. 45, 836–839.

21. Lanin, A., Chebotarev, A., Kelmanson, I., Pochechuev, M., Fetisova, E., Bilan, D., Shevchenko, E., Ivanov, A., Fedotov, A., and Belousov, V. (2021). Single-beam multimodal nonlinear-optical imaging of structurally complex events in cell-cycle dynamics. JPhys. Photonics 3, 044001.

22. Sun, C.K., Kao, C.T., Wei, M.L., Chia, S.H., Kärtner, F.X., Ivanov, A., and Liao, Y.H. (2019). Slide-free imaging of hematoxylin-eosin stained whole-mount tissues using combined third-harmonic generation and three-photon fluorescence microscopy. J. Biophotonics 12, e201800341.

23. Chu, S.W., Tai, S.P., Ho, C.L., Lin, C.H., and Sun, C.K. (2005). High-resolution simultaneous three-photon fluorescence and third-harmonic-generation microscopy. Microsc. Res. Tech. 66, 193–197.

24. Wang, C., and Yeh, A.T. (2012). Two-photon excited fluorescence enhancement with broadband versus tunable femtosecond laser pulse excitation. J. Biomed. Opt. 17, 025003.

25. Studier, H., Breunig, H.G., and König, K. (2011). Comparison of broadband and ultrabroadband pulses at MHz and GHz pulse-repetition rates for nonlinear femtosecond-laser scanning microscopy. J. Biophotonics 4, 84–91.

26. Chudoba, C., Fujimoto, J., Ippen, E., Haus, H., Morgner, U., Kärtner, F., Scheuer, V., Angelow, G., and Tschudi, T. (2001). All-solid-state Cr: forsterite laser generating 14-fs pulses at 1.3 μm. Opt. Lett. 26, 292–294.

27. Chia, S.-H., Liu, T.-M., Ivanov, A.A., Fedotov, A.B., Zheltikov, A.M., Tsai, M.-R., Chan, M.-C., Yu, C.-H., and Sun, C.-K. (2010). A sub-100fs self-starting Cr: forsterite laser generating 1.4 W output power. Opt. Express 18, 24085–24091.

28. Ivanov, A., Minkov, B., Jonusauskas, G., Oberle, J., and Rulliere, C. (1995). Influence of Cr4+ ion concentration on cw operation of forsterite laser and its relation to thermal problems. Opt. Commun. 116, 131–135.

29. Berlier, J.E., Rothe, A., Buller, G., Bradford, J., Gray, D.R., Filanoski, B.J., Telford, W.G., Yue, S., Liu, J., and Cheung, C.-Y. (2003). Quantitative comparison of long-wavelength Alexa Fluor dyes to Cy dyes: fluorescence of the dyes and their bioconjugates. J. Histochem. Cytochem. 51, 1699–1712.

30. Akbari, N., Rebec, M.R., Xia, F., and Xu, C. (2022). Imaging deeper than the transport mean free path with multiphoton microscopy. Biomed. Opt. Express 13, 452–463.

31. Tao, X., Lin, H.-H., Lam, T., Rodriguez, R., Wang, J.W., and Kubby, J. (2017). Transcutical imaging with cellular and subcellular resolution. Biomed. Opt. Express 8, 1277–1289.

32. Campusano, J.M., Su, H., Jiang, S.A., Sicaeros, B., and O’Dowd, D.K. (2007). nAChR-mediated calcium responses and plasticity in Drosophila Kenyon cells. Dev. Neurobiol. 67, 1520–1532.

33. Lyutova, R., Selcho, M., Pfeuffer, M., Segebarth, D., Habenstein, J., Rohwedder, A., Frantzmann, F., Wegener, C., Thum, A.S., and Pauls, D. (2019). Reward signaling in a recurrent circuit of dopaminergic neurons and peptidergic Kenyon cells. Nat. Commun. 10, 1–14.

34. Sun, X.R., Badura, A., Pacheco, D.A., Lynch, L.A., Schneider, E.R., Taylor, M.P., Hogue, I.B., Enquist, L.W., Murthy, M., and Wang, S.S.-H. (2013). Fast GCaMPs for improved tracking of neuronal activity. Nat. Commun. 4, 1–10.

